# A machine learning model for screening of body fluid cytology smears

**DOI:** 10.1101/2021.07.20.453010

**Authors:** Parikshit Sanyal, Sayak Paul, Vandana Rana, Kanchan Kulhari

## Abstract

**Introduction:** Body fluid cytology is one of the commonest investigations performed in indoor patients, both for diagnosis of suspected carcinoma as well as staging of known carcinoma. Carcinoma is diagnosed in body fluids by the pathologist through microscopic examination and searching for malignant epithelial cell clusters. The process of screening body fluid smears is a time consuming and error prone process.

**Aim:** We have attempted to construct a machine learning model which can screen body fluid cytology smears for malignant cells.

**Materials and methods:** MGG stained Ascitic / pleural fluid cytology smears were included from 21 cases (14 malignant, 07 benign) in this study. A total of 693 microphotographs were taken at 40x magnification at the same illumination and after correction of white balance. A Magnus Microphotography system was used for photography. The images were split into the training set (195 images), test set (120 images) and validation set (378 images).

A machine learning model, a convolutional neural network, was developed in the Python programming language using the Keras deep learning library. The model was trained with the images of the training set. After completion of training, the model was evaluated on the test set of images.

**Results:** Evaluation of the model on the test set produced a sensitivity of 97.87%, specificity 85.26%, PPV 95.18%, NPV 93.10% In 06 images, the model has failed to detect singly scattered malignant cells/ clusters. 14 (3.7%) false positives was reported by the model.

The machine learning model shows potential utility as a screening tool. However, it needs improvement in detecting singly scattered malignant cells and filtering inflammatory infiltrate.

## Introduction

Ascitic fluid cytology is one of the first tests conducted in a patient with ascites, both for confirmation of a suspected malignancy and staging of a known malignancy. The detection of malignant cells in ascitic fluid is carried out by the pathologist by light microscopic examination. The sensitivity of this method in in detecting malignancy has been found to be 50% – 96.7% in different studies.^[1] [2] [3] [4]^Qne large case series found that ovarian malignancy is the only malignancy yielding a significant rate of detection from ascitic fluid cytology. ^[1]^ There is a significant proportion of cases which are reported either false positive of false negative in ascitic fluid cytology. In one study of 170 patients, peritoneal cytology showed false negative results in 30.02% cases, while in 6.38% the results were false positive.^[3]^ This large variability in accuracy may be attributable to the often sparse and uneven distribution of malignant cell clusters in the smear. Thus screening slides for malignant cells is often a time consuming and error prone process. In one study, the false-negative were attributed to sampling errors in 71% of pleural and 73% of peritoneal effusions and to screening errors in 29% and 27%, respectively.^[4]^ Immunostains for epithelial markers such as B72.3, MOC-31, and Ber-EP4 have been suggested for easy identification of such malignant clusters and also to confirm their epithelial origin.^[5]^ Preparation of cell blocks add 10-15% to the diagnostic sensitivity of body fluid cytology. ^[6][7].^ . In addition, immunohistochemical markers such as CEA significantly enhance the diagnostic efficacy: a combination of calretinin negative and CEA positive staining showed 97% sensitivity and 100% specificity for malignancy in one study of 50 cell blocks.^[8]^ But in a large case series of 5.5 years, it was also shown that use of IHC markers produces a higher rate of indeterminate but not malignant diagnosis ^[9]^, thus undermining the utility of IHC.

Artificial neural networks (ANNs) are a large family of trainable models, where each subfamily of models is optimized for different functions. We have chose the ANN subfamily, known as convolutional neural networks (CNNs) which are shown to perform image-based object classification.^[10]^ CNNs take a whole image as input and classifies the image in defined categories. The input image is passed through multiple “layers”, each layer comprising multiple linear convolutional filters.^[11]^ The input for each layer is the output of the previous one, with an overlaid non-linearity. The image “features” extracted by the layers are finally fed into a classifier that determines the category the image belongs to. CNNs have been described by Karpathy et al.^[12]^

Deep, multilayered CNNs have been successful in recognising everyday objects^[11,13]^ and classifying them in correct categorie^[14]^ We have aimed to apply the principles of CNNs to identify malignant cell clusters from ascitic fluid cytology smears. The model should be able to correctly identify malignant cell clusters from microphotographs of ascitic fluid cytology smears, and present them to the pathologist for review. The proportion of false negatives and false positives will have to be kept at a minimum. The model would serve as a pathologist assistant and a screening tool, and present foci of clusters identified as ‘malignant’ to the pathologist for review.

## Materials and methods

Cases were selected from the archives of two hospitals. May Grunwald Giemsa stained Ascitic fluid/ pleural fluid cytology smears were included from 21 cases (14 malignant, 07 benign) in this study. All 14 malignant cases were confirmed by histopathology (except the one case of carcinoma of unknown primary site) . The presence of malignant epithelial cells in ascitic/ pleural fluid were confirmed independently by two pathologists.

A total of 693 microphotographs were captured at 40x magnification at the same illumination and after correction of white balance. The foci were manually chosen as to represent well defined benign or malignant cells, and reviewed by two independent pathologists. A Magnus Integrated Microphotography system was used for photography. The photographed images were of 1280 × 960 pixels in resolution.

The images were split into two subsets (Table 1).

1. a training set – for training of the machine learning model (i.e. learn the points of difference between appearance of benign and malignant cells in cytology)
2. a test set – for concurrent performance evaluation during training
3. a validation set – to evaluate the performance of the model after completion of training

**Table 1:**
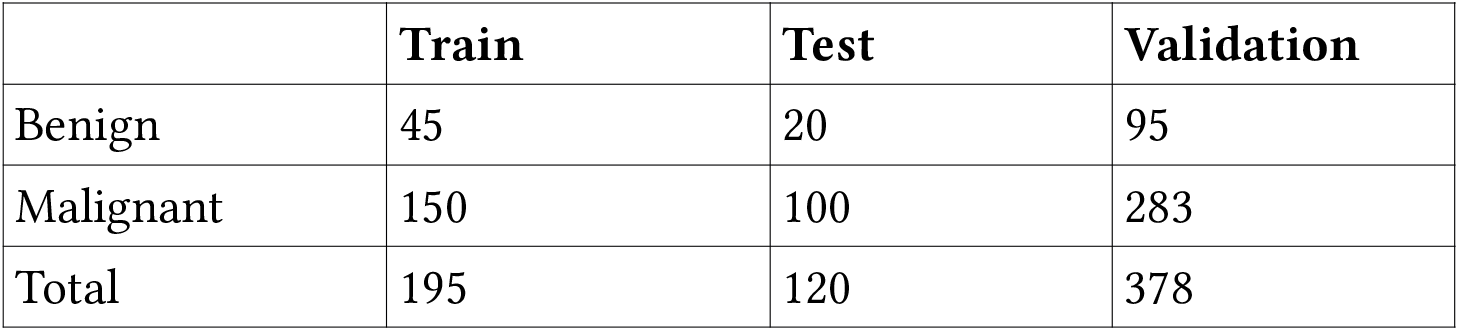
Distribution of images in training, test and validation set.

|  | **Train** | **Test** | **Validation** |
| --- | --- | --- | --- |
| Benign | 45 | 20 | 95 |
| Malignant | 150 | 100 | 283 |
| Total | 195 | 120 | 378 |

The machine learning model was developed in the Python programming language using the Keras deep learning library. A multilayered deep convolutional neural network (CNN) was developed using successive layers of convolutions, rectified linear unit (ReLu) and fully connected layers. The CNN takes an color image 256 × 192 pixels in dimension, and produces an output ‘0’ (corresponding to benign) and ‘1’ (malignant). The architecture of the CNN is shown in Figure 1.

**Figure 1:**
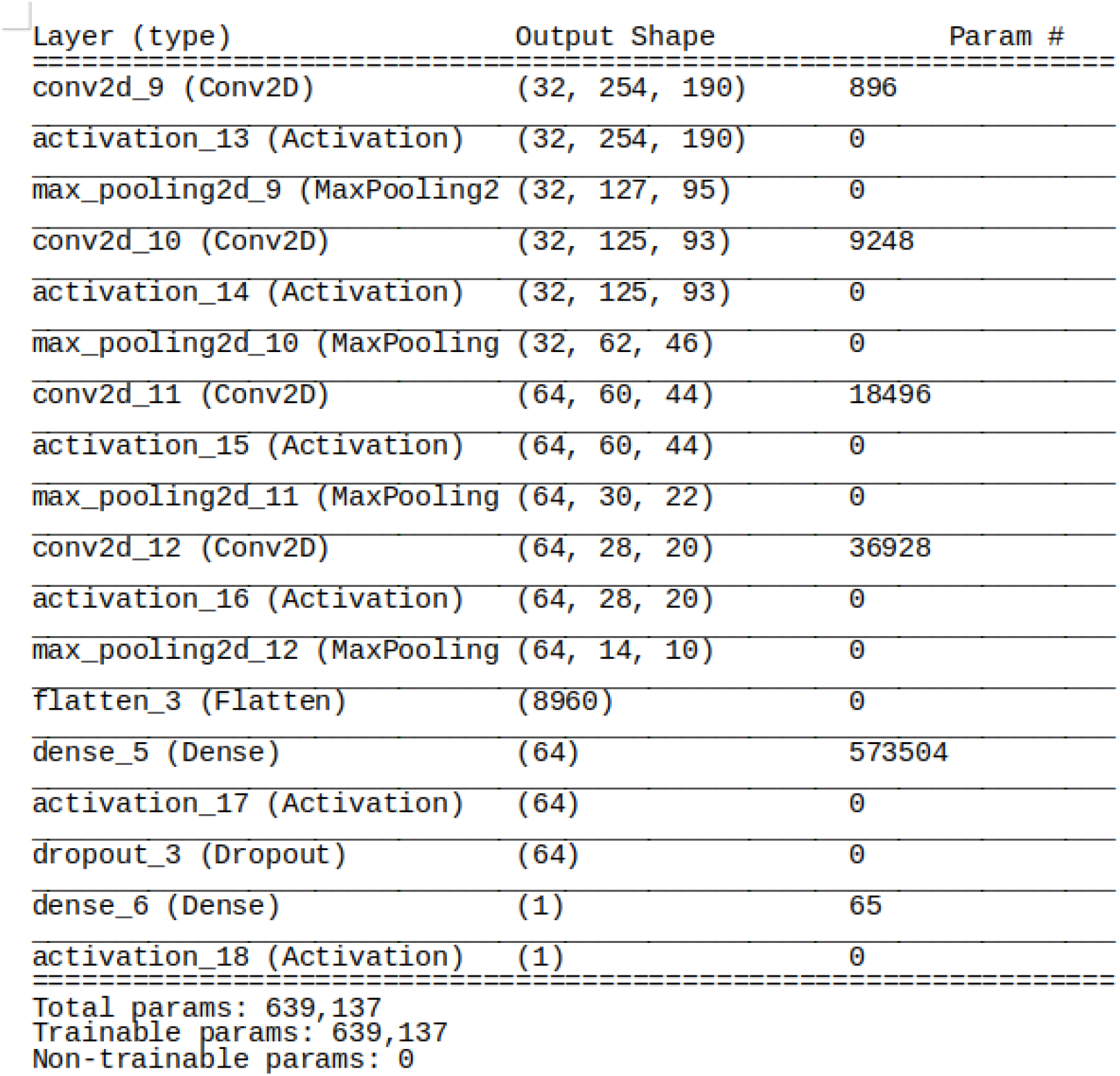
Architecture of the Convolutional Neural Network.

Images were resized to 256 × 192 pixels before training. The CNN was trained several times with the images of the training set, with varying number of epochs; an optimal training with 10 epochs of and 200 steps per epoch was chosen (Figure 2) which maximised the diagnostic accuracy. Over the period of training, the CNN adjusted its parameters to produce the correct output in majority of cases. During training, 93.75% accuracy was achieved on the testing set. The loss function (error rate) showed gradual decline over both training and test set. (Figure 3–4)

**Figure 2:**
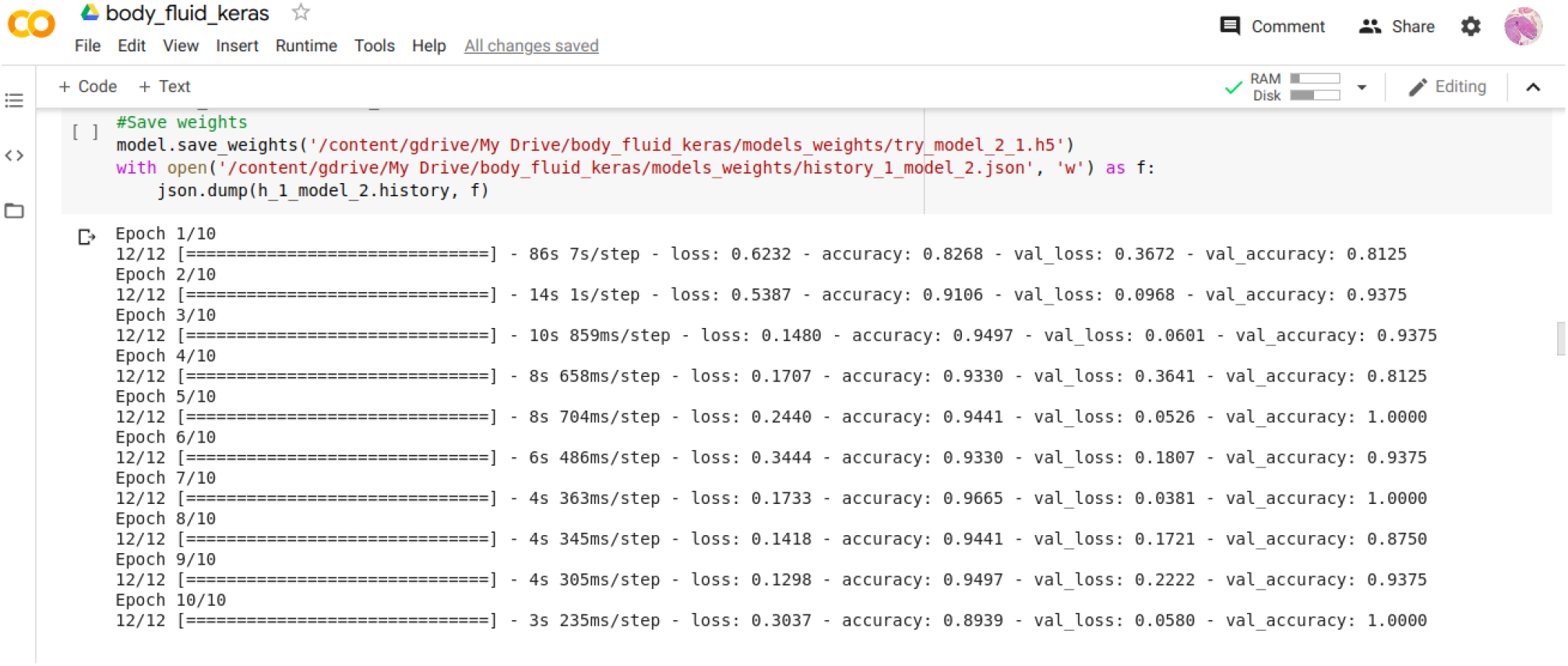
Training session of the model.

**Figure 3:**
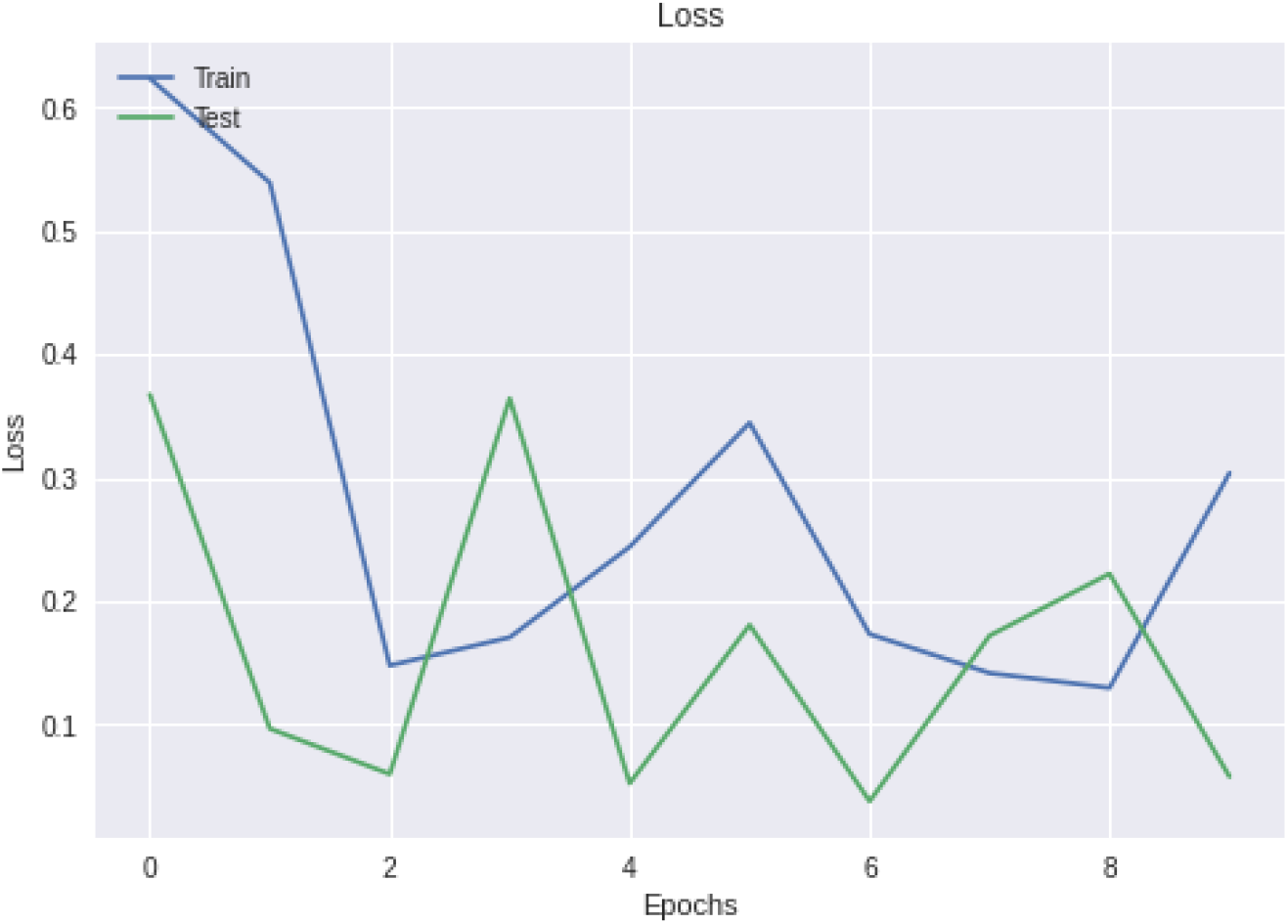
Loss function (error rate) plotted versus epochs of training.

**Figure 4:**
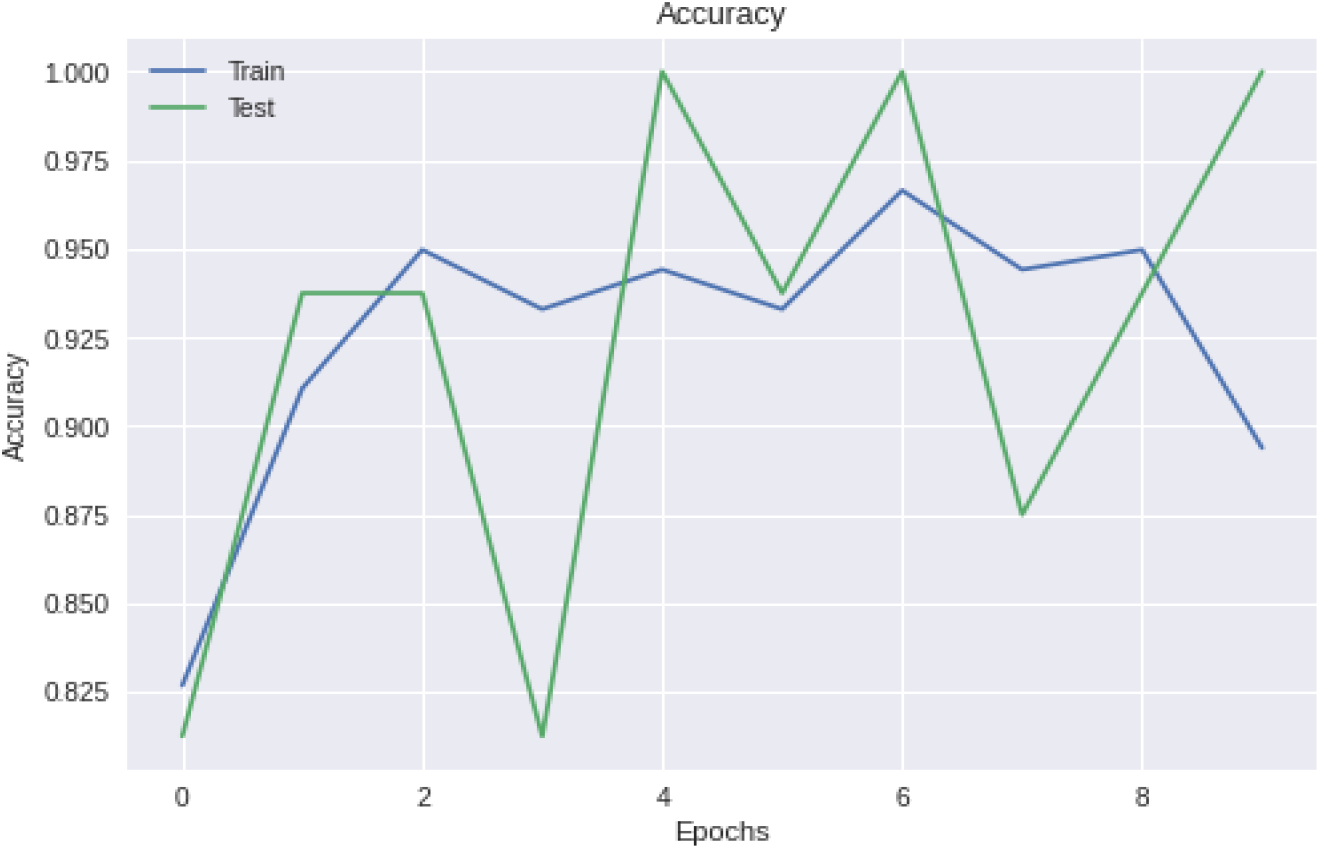
Accuracy of the model plotted versus epochs of training.

After completion of training, the performance of the model on the validation set of images was evaluated using standard statistical methods.

## Results

Evaluation of the model on the test set produced 97.87% sensitivity, 85.26% specificity, 95.18% positive predictive value (PPV) and 93.10% negative predictive value (NPV). The model successfully classified majority of benign foci. Only 06 false negatives and 14 false positive was reported (Table 2).

**Table 2:** Performance of the model on the validation set.

|  | Actual |  |  |
| --- | --- | --- | --- |
|  | Benign | Malignant | Total |
| Predicted |  |  |  |
| Benign | 81 | 6 | 87 |
| Malignant | 14 | 277 | 291 |
| Total | 95 | 283 | 378 |
| Sensitivity | 97.87% | Specificity | 85.26% |
| PPV | 95.18% | NPV | 93.10% |

In 06 (1.5%) images, the model has failed to detect malignant cells; all of these images were paucicellular. Possible over-fitting has resulted in 14 false positive images; in 11 of these images, a hemorrhagic background seems to have interfered with correct classification. (Figure 5–8)

**Figure 5:**
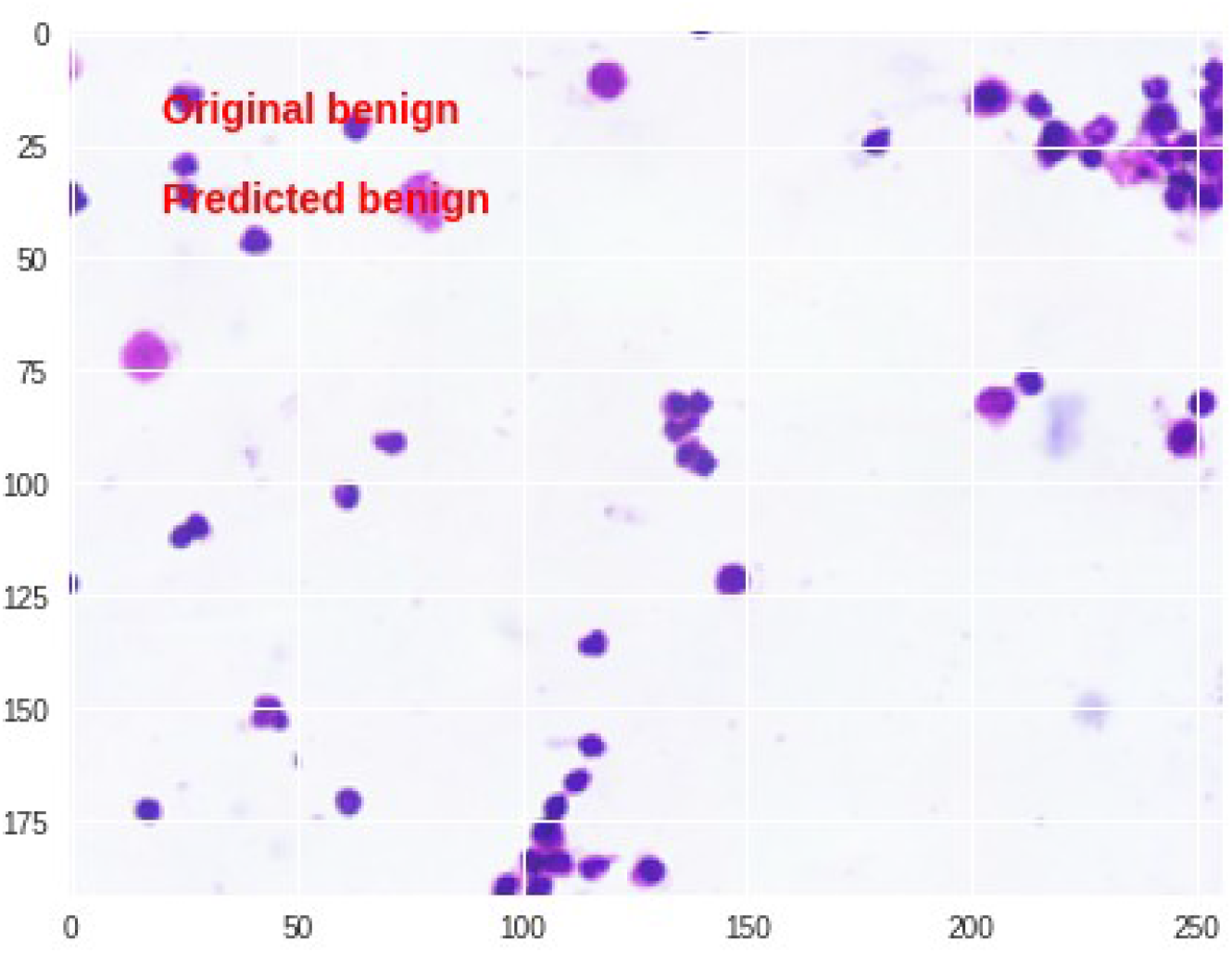
Correct classification by the model.

**Figure 6:**
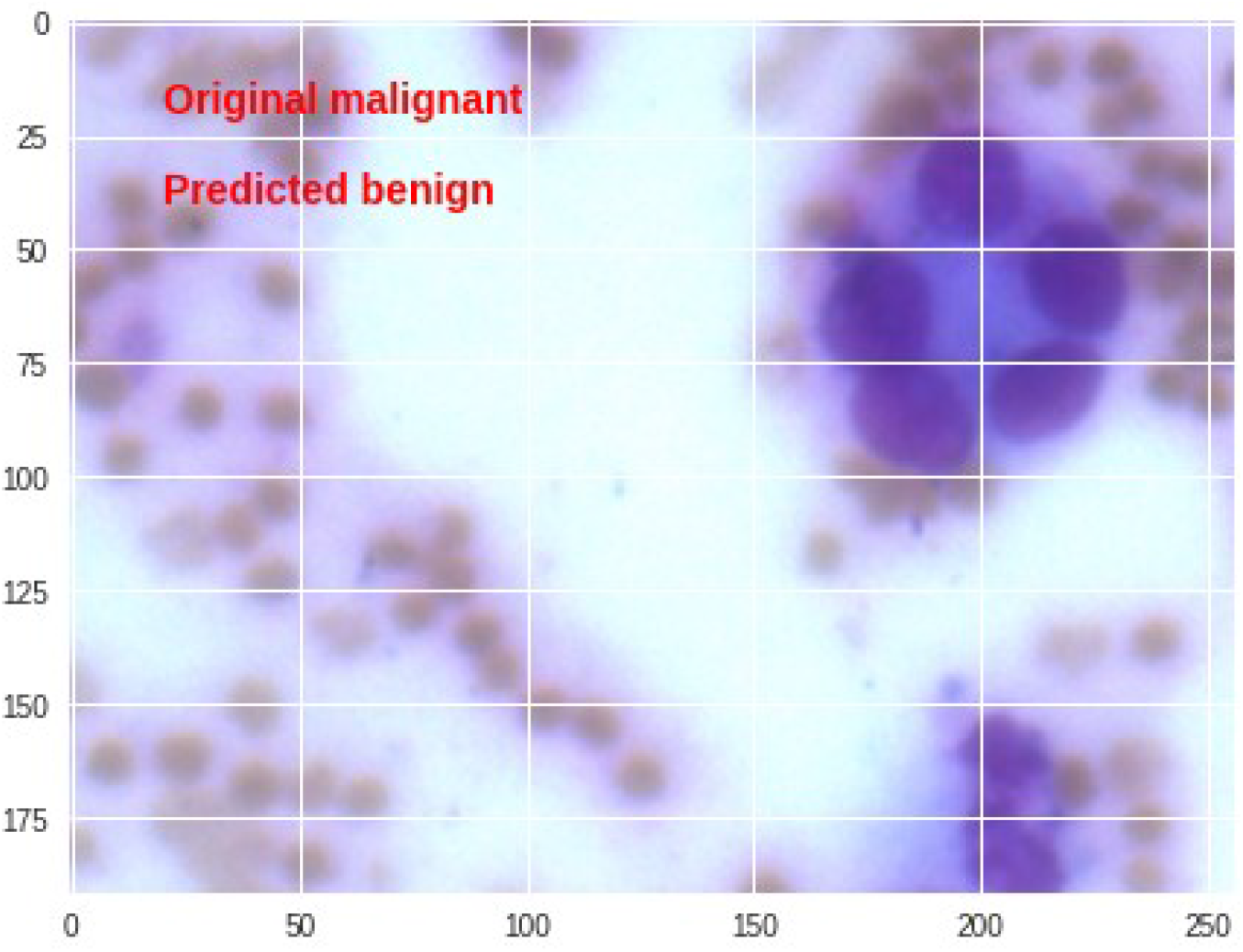
The model has missed the single cluster of malignant cells.

**Figure 7:**
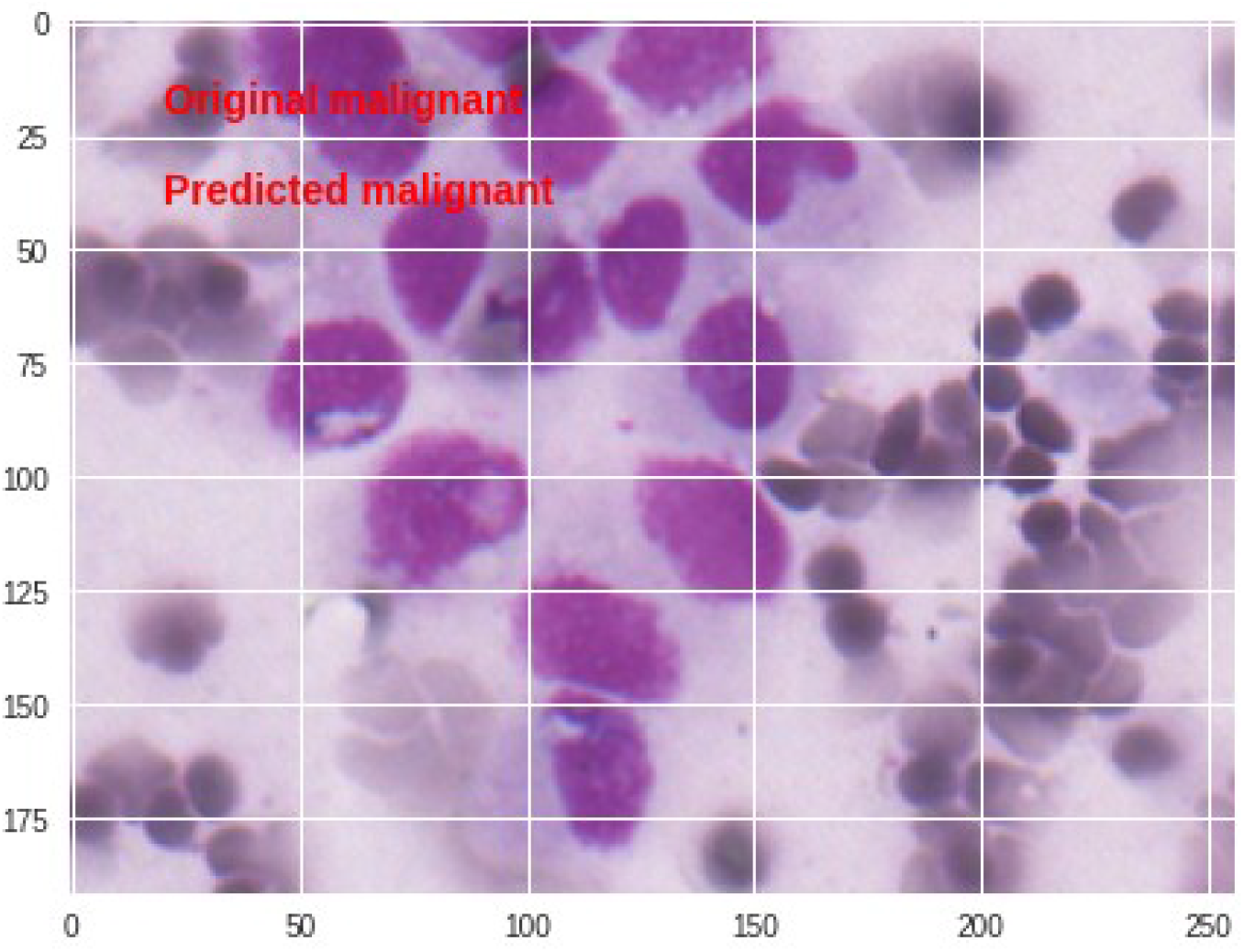
Malignant cell cluster correctly identified by the model.

**Figure 8:**
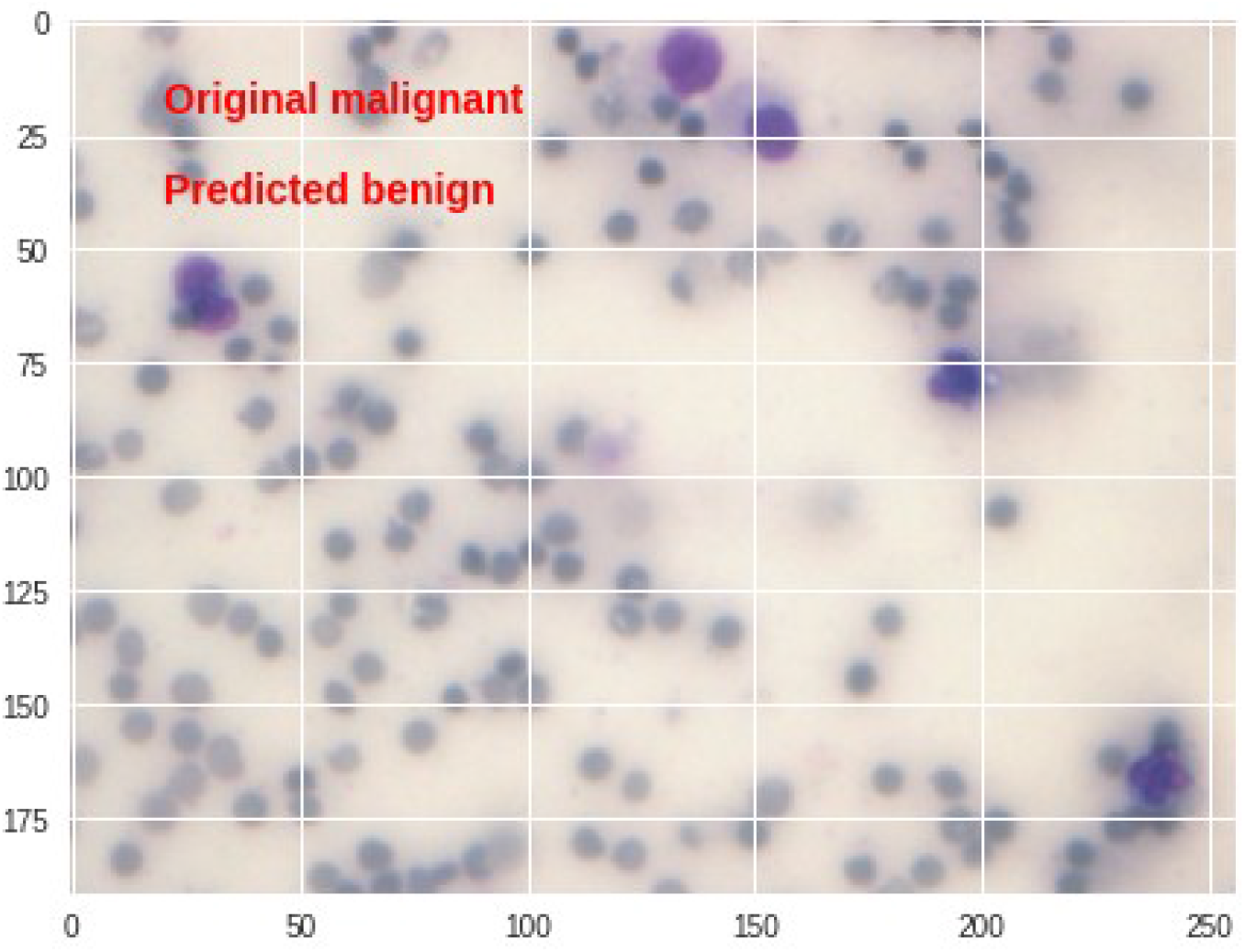
Paucity of cells have interfered with the identification of malignant cells in this focus.

The results demonstrate that the model is capable of correctly classifying majority of foci in benign or malignant categories. A rare mesothelial cell cluster has been labeled malignant (Figure 9). The model has demonstrated the ability to detect malignant cells from a densely inflammatory background (Figure 10). However, difficulties remain where cellularity is low and malignant cells are present in small isolated clusters. Also, a hemorrhagic background can obscure malignant cells.

**Figure 9:**
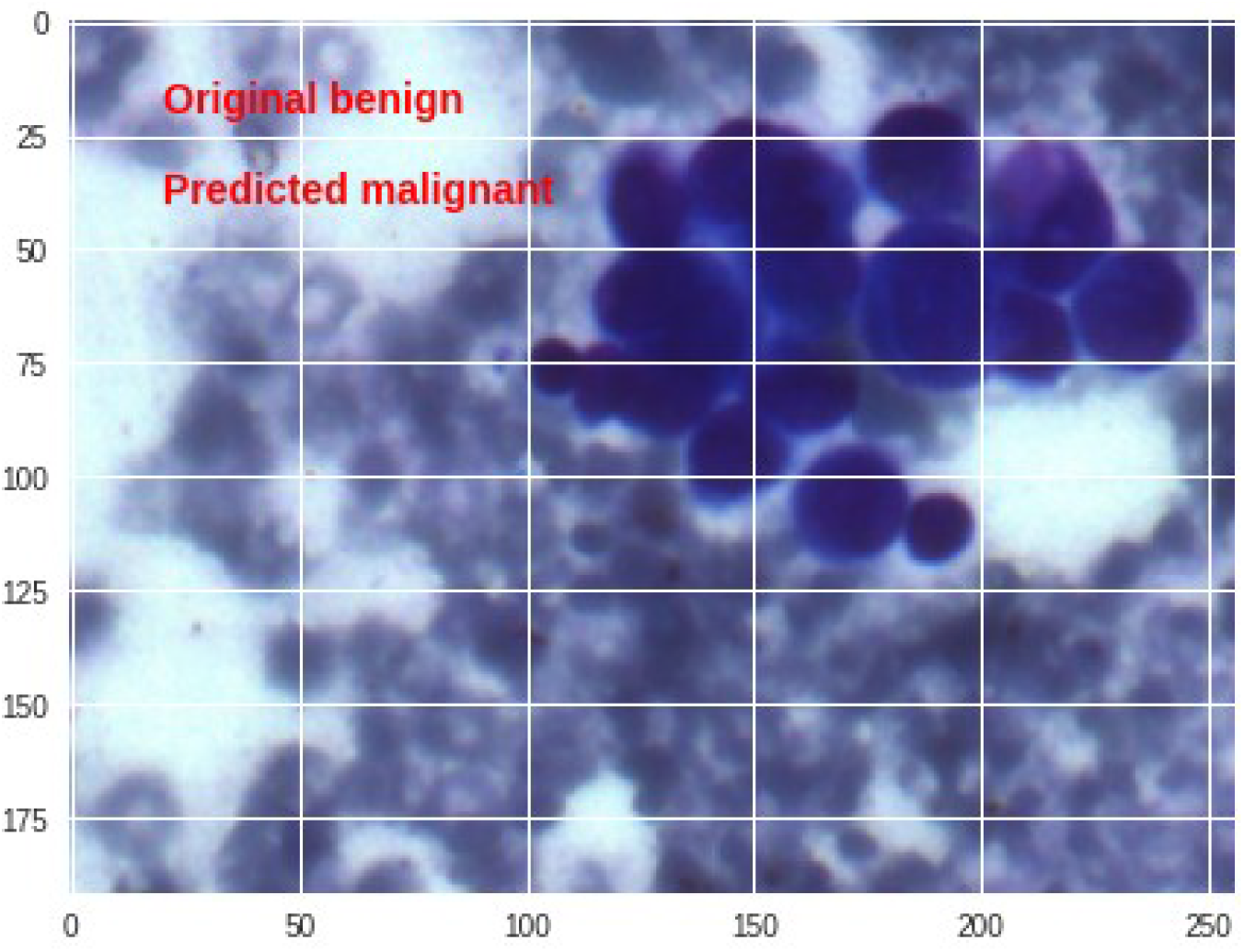
Mesothelial cell clusters wrongly classified as malignant, over a hemorrhagic background.

**Figure 10:**
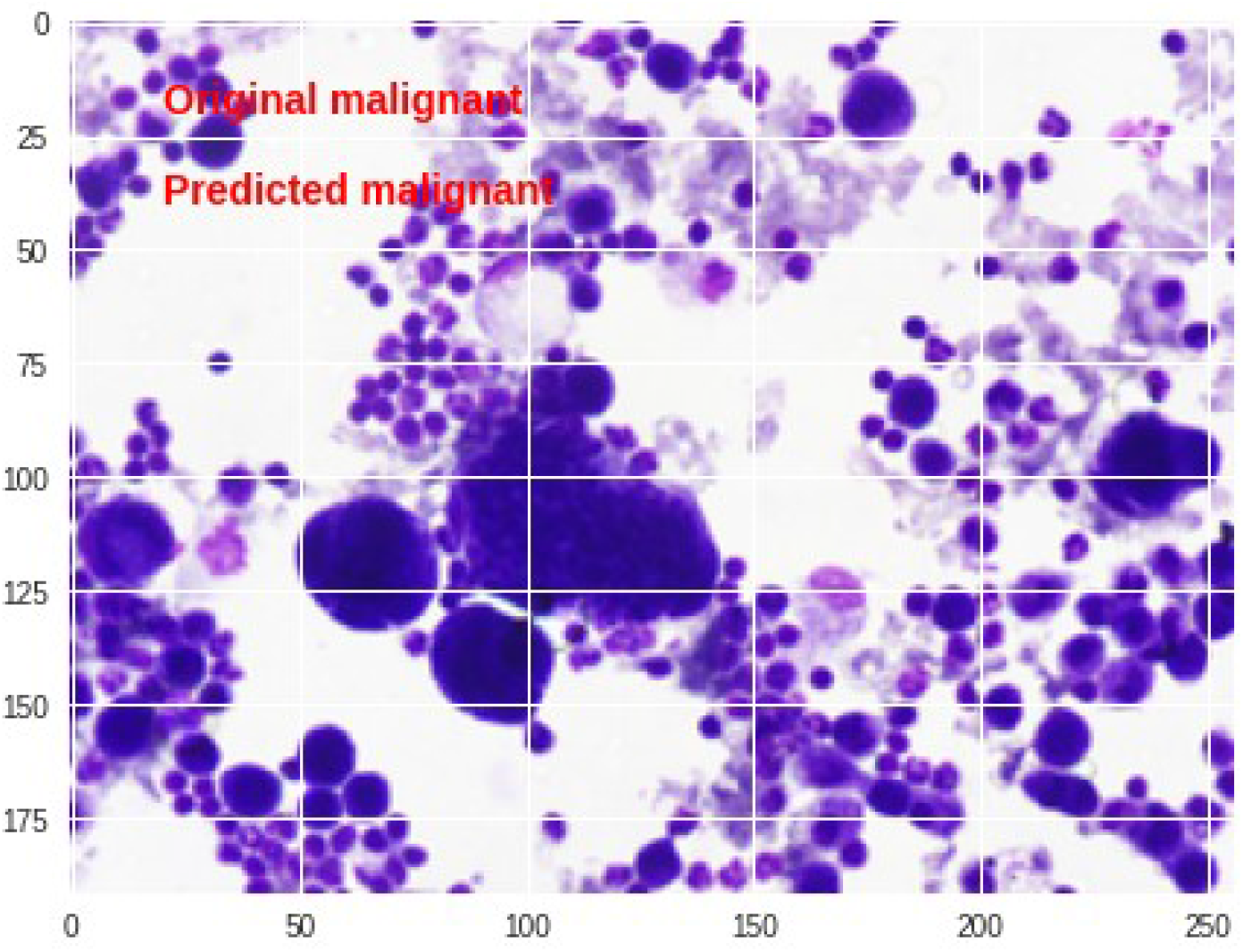
Malignant cells detected over an inflammatory background.

Analysis of receiver operating characteristics of the model showed area under curve (AUC) to be 0.916, which is the diagnostic accuracy of the model (Figure 11).

**Figure 11:**
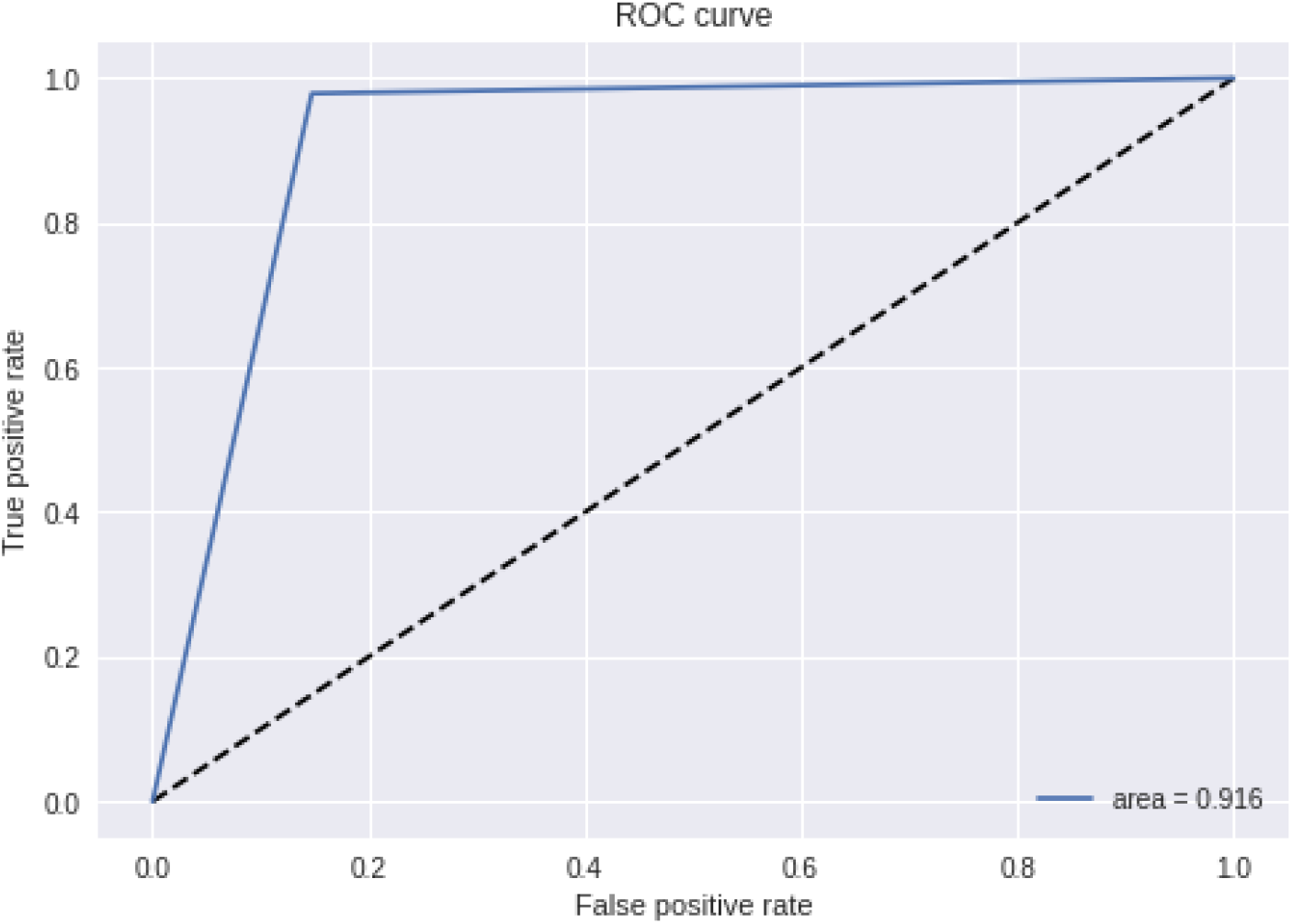
Receiver operating characteristics (ROC) of the model.

## Discussion

Whereas microscopy on direct MGG stained smears is the traditionally accepted method, it suffers from the drawbacks of low sensitivity and variable inter-observer reproducibility. Cell block preparation and IHC have been found to be of optimal value in only 10-12% of fluids (in a population with relatively low prevalence of malignancy), above which their diagnostic utility diminishes.^[9]^ The sensitivity and specificity of IHC in detection of malignancy in body fluid has yet not exceeded 90%.^[20] [21]^

Artificial neural networks have emerged as a useful decision support tool in cytopathology. ^[15]^ Pathologist’s assistant machine learning models have been developed for cervical^[16] [17]^ and thyroid ^[18] [19]^ cytology. The present study aims to develop such a model for recognising malignant cells from body fluid cytology smears.

There have been few studies regarding such automated decision systems on body fluid cytology. There has been focus in extracting geometric features from the image and applying machine learning models over the extracted features. For example, Win et al used extracted features from the image and used them as inputs for an artificial neural network. On a sample of 125 images, their method achieved sensitivity of 87.97%, specificity of 99.40%, with 98.70% diagnostic accuracy. ^[22]^ Baykal et al used the technique of active appearance model for to achieve effective cell segmentation from cytopathological images, with good diagnostic accuracy.^[23]^ The wavelet transform has also been shown to achieve a high recognition ratio. ^[24]^ Zhang et al have used morphometric parameters (area rate of the karyon and cytoplasm, the optic density, the shape factor) and used these parameters with a fuzzy pattern recognition model to detect cancer cells. ^[25]^ In a related study, while analysing the difference between malignant and benign mesothelial cell proliferations, Tosun et al found that the quantification of chromatin distribution is 100% predictive of whether a cell is malignant.^[26]^

We have not extracted any geometric feature from images, because we have included images from a variety of foci from which such morphometry is not possible. Our study sample has included smears with eosinophilic material rich background, few foci with dense inflammatory infiltrate, and also few foci with very sparse malignant cells. Keeping all this variation in mind, we have used the entire image as input to the neural network. This method, however, has yielded 97% sensitivity (on individual foci), albeit with moderate specificity (85%).

A subset of the intermediate layers of the neural network as it processes an image is shown in Figure 12. The image is simplified and filtered over the layers of the network, to finally produce an output of 0 or 1. It is interesting to note how a certain future of an image gets highlighted and extracted over several convolutions and transformations through the layers of the network.

**Figure 12:**
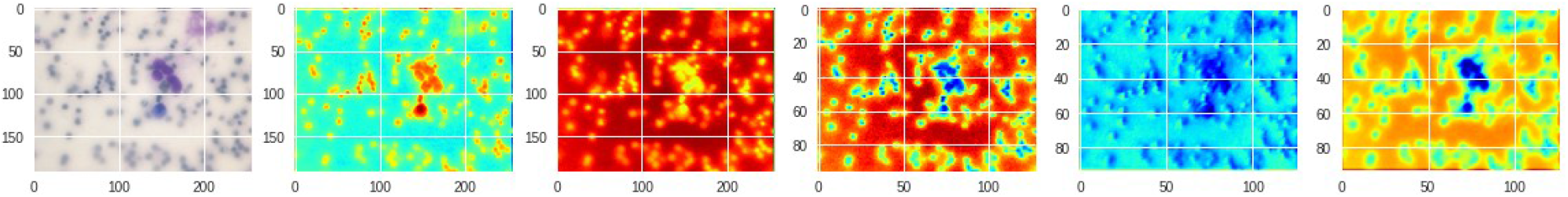
Processing an image through the intermediate layers of the CNN.

It is important to recognize the limitations of this model: hemorrhagic background has been shown to interfere with its functioning (Figure 9) , and it has failed to recognize two foci of sparse malignant cells (Figure 10). The study was performed with a limited number of samples to develop a prototype machine learning model. Further refinement in technique and training with a greater number of images are required.

## Conflicts of interest

None to declare

## Data availability

Image dataset and code are available at https://github.com/vaishleshik/body_fluid_classifier.

## Fundings

No external funding was received for this study

## Note

Sayak Paul made his contribution while employed at PylmageSearch

## Notes

https://github.com/vaishleshik/body_fluid_classifier

### Competing Interest Statement

The authors have declared no competing interest.

### Summary of Updates

The author order has been modified to match the uploaded PDF

